# Genomic diversity, antibiotic resistance, and maturation‑dependent adhesion of F18 enterotoxigenic *Escherichia coli* strains in porcine intestinal cells

**DOI:** 10.64898/2026.01.23.701439

**Authors:** Dongqi Liu, Nicholas L.F. Gallina, Chenhai Li, Nicole Irizarry-Tardi, Mahmoud Sayedahmed, Jathya C. Karunathilaka, Nathan Horn, Weicang Wang, Ahmed Abdel Khalek, Arun K. Bhunia

## Abstract

Enterotoxigenic *Escherichia coli* (ETEC) strains expressing F4 and F18 fimbriae are major causes of neonatal and post-weaning diarrhea in swine. Although epithelial maturation influences susceptibility in vivo, its impact on ETEC-host interactions remains poorly defined. This study characterizes emerging F18 ETEC isolates using a differentiated porcine intestinal cell model. Three F18 strains (3EC1, 27EC1, 3247EC), a porcine F4 strain, and human ETEC H10407 were analyzed by comparative genomics for virulence factors, toxin genes, and antimicrobial resistance determinants. Adhesion assays were performed using IPEC-1, IPEC-J2, and Caco-2 cells conditioned to Early (6 days post-confluence, DPC), Mid (9 DPC), and Late (16 DPC) maturation states. Transcription of F18-binding receptors (FUT1, FUT2) was quantified by RT-qPCR. IPEC-1 cells exhibited significantly higher FUT1 and FUT2 expression than IPEC-J2, corresponding to approximately two-fold stronger adhesion by most F18 isolates. Strain 3EC1 showed a distinct adhesion peak at 9 DPC, approaching F4 levels, while F4 and H10407 consistently displayed the highest adhesion across all models. Genomic analyses revealed substantial heterogeneity among F18 strains in fimbrial loci, flagellin, lipopolysaccharide biosynthesis, and antimicrobial resistance. Strain 3EC1 uniquely carried *stx2e*, and non-classical EAST1 variants were detected in 3EC1 and 3247EC. All F18 isolates encoded *hlyE* and were β-hemolytic; 3247EC harbored 28 antimicrobial resistance genes. The IPEC-1/IPEC-J2 maturation stages recapitulate age-dependent susceptibility to ETEC, likely driven by FUT1/2 expression levels. The combination of strong adhesion, *stx2e*, and extensive antimicrobial resistance in F18 strains underscores their evolving virulence and supports this model as a refined platform for studying porcine ETEC pathogenesis.

**Importance:** ETEC remains a leading cause of neonatal and post-weaning diarrhea in swine, yet the biological basis for age-dependent susceptibility is not fully understood. This study demonstrates that maturation of porcine intestinal epithelial cells strongly influences F18 ETEC adhesion, driven in part by developmental regulation of the F18-binding receptors *FUT1* and *FUT2*. By integrating comparative genomics with a physiologically relevant *in vitro* maturation model, we reveal substantial diversity in virulence and resistance among F18 strains, including strong adhesion capacity, *stx2e*, and extensive antimicrobial resistance in strain 3EC1. These findings highlight the evolution of ETEC toward increased persistence and pathogenic potential in swine populations. The interaction between the maturation-dependent IPEC-1 and IPEC-J2 cell lines and ETEC offers a valuable tool for evaluating intervention strategies to reduce weaning piglet susceptibility to ETEC infection.

## Introduction

Enterotoxigenic *Escherichia coli* (ETEC) is a leading bacterial pathogen responsible for piglet diarrhea and edema disease (ED) (1,2) and contributes to significant weight loss, high morbidity, and mortality. Post-weaning diarrhea (PWD), a hallmark of ETEC infection, imposes substantial financial burdens on the swine industry due to reduced growth rates and increased veterinary costs (3).

Current control strategies, including zinc supplementation, antibiotics, probiotics, and prebiotics, are increasingly challenged by the emergence of antimicrobial resistance (AMR), driven by the overuse of antibiotics in livestock feed (4-6), particularly in the United States (7,8). These challenges underscore the urgent need to elucidate the ETEC virulence mechanism to develop sustainable alternatives to antibiotics.

Fimbriae are a critical virulence factor that mediates ETEC adhesion and colonization in the porcine small intestine (9). ETEC expressing F4 (K88) or F18 (F107, 2134P, 8813) fimbriae exhibits distinct age-specific pathogenicity: F4 predominantly affects neonate piglets, while F18 is associated with PWD in weaned piglets (7,10). F4 fimbriae are categorized into three subtypes (K88ab, K88ac, K88ad), while F18 fimbriae are divided into F18ab (associated with ED) and F18ac (associated with PWD) (11). These differences in tropism are attributed to the dynamic expression of host intestinal receptors. F4 receptors (e.g., MUC4) are highly expressed after birth and start to decline after weaning (12,13), whereas F18 receptors (e.g., FUT1) increase in expression at three weeks of age and persist into adulthood (14). However, conflicting evidence suggests comparable FUT1 mRNA levels in newborn and weaned piglets (15), highlighting unresolved questions about F18 receptor dynamics. The F18 operon comprises five genes (*fedA*–*fedF*), with FedA forming the structural backbone. FedE and FedF mediate receptor binding alongside FedA (16). FedA and FedB also facilitate pilus assembly (17).

This study characterized three disease-causing *E. coli* strains isolated from swine farms through biochemical, genetic, and functional assays. We utilized porcine (IPEC-1 and IPEC-J2) (18-21) and human intestinal epithelial (Caco-2) cell lines with differential susceptibility to F4- and F18-pathotype ETEC to compare and evaluate their virulence. Two porcine cell lines were assessed at varying stages of maturity to investigate how intestinal development influences bacterial adhesion and pathogenicity. Whole-genome sequencing was applied to interrogate the genetic profiles of these isolates, shedding light on the novel acquisition of virulent features and antibiotic resistance in the current epidemiology. The antibiotic resistance profile of each strain was also examined. Our findings aim to advance understanding of ETEC evolution, host-pathogen interactions, and antibiotic resistance to inform the development of targeted interventions.

## RESULTS

### Virulence factors characterization in ETEC F4 and ETEC-F18

ETEC cultures (3EC1, 27EC1, and 3247EC) used in the study were further verified to be lactose fermenting, showing growth on MacConkey agar (**Fig. S1**) and hemolytic (**Fig. 1A**). PCR confirmed the presence of the *fedA* (506 bp) in all F18 *E. coli* isolates (3EC1, 27EC1, and 3247EC) since *fedA* is a distinctive marker correlated with PWD or edema disease (22) and encodes the core structure of F107 fimbriae used to classify the ETEC-F18 strains (11,23). Non-ETEC F18 strains, such as ETEC F4 (K88), ETEC F5 (K99), and *Listeria monocytogenes* F4244 (*Lm*), used as controls, tested negative for *fedA* (**Fig. 1B**).

**FIG 1.**
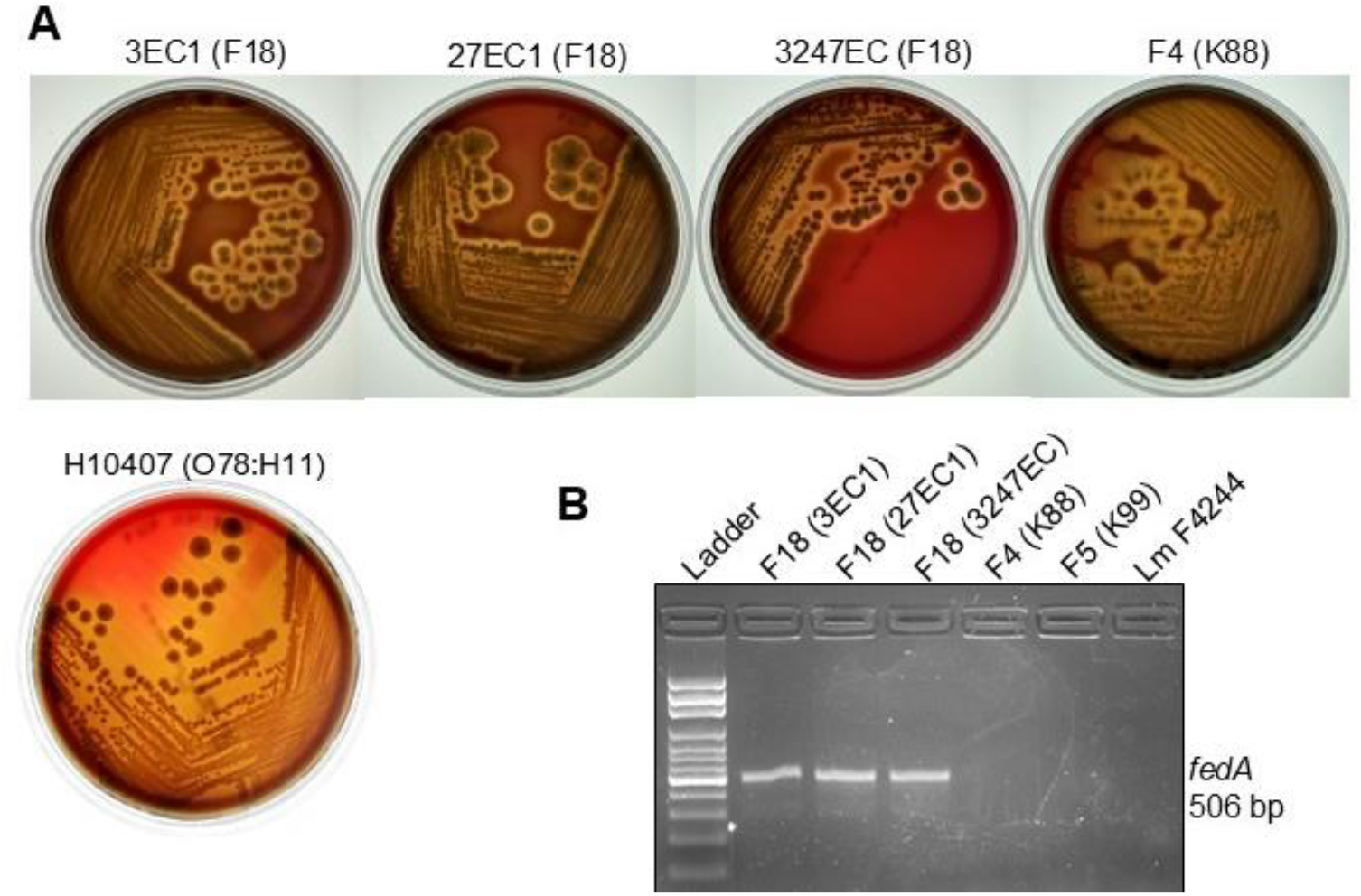
Characterization of Enterotoxigenic *Escherichia coli* (ETEC) F18 strains. (**A**) Hemolytic activity assay on sheep-red blood agar plates showing clear hemolytic zones surrounding the colonies in all strains tested. (**B**) PCR confirmation of enterotoxigenic *Escherichia coli* (ETEC) F18 strains showing amplification of *fedA* gene (506 bp), while ETEC F4 and F5 and *Listeria monocytogenes* (Lm) F4244 showed no amplification as negative controls.

### Age-differentiated adhesion comparison of ETEC F4 and F18 on IPEC-J2

The maturation of intestinal epithelial cells influences the relative expression of ETEC fimbriae-specific receptors (13,14,18). To investigate this, IPEC-1 and IPEC-J2 cells were categorized into three distinct maturity phases: “Early” (6 days post-confluence, DPC), “Mid” (9 DPC), and “Late” (16 DPC) (**Fig. 2**). In the “Early” phase, the cell monolayer displayed a clear epithelioid morphology with well-defined cellular boundaries. As differentiation progressed, cellular morphology became increasingly complex. By the “Mid” phase, intercellular overlap became apparent, and cellular boundaries appeared less distinct. At the “Late” phase, notable cytoplasmic changes were observed, including granule accumulation and vacuolation (**Fig. 2A** and **C**).

**FIG 2.**
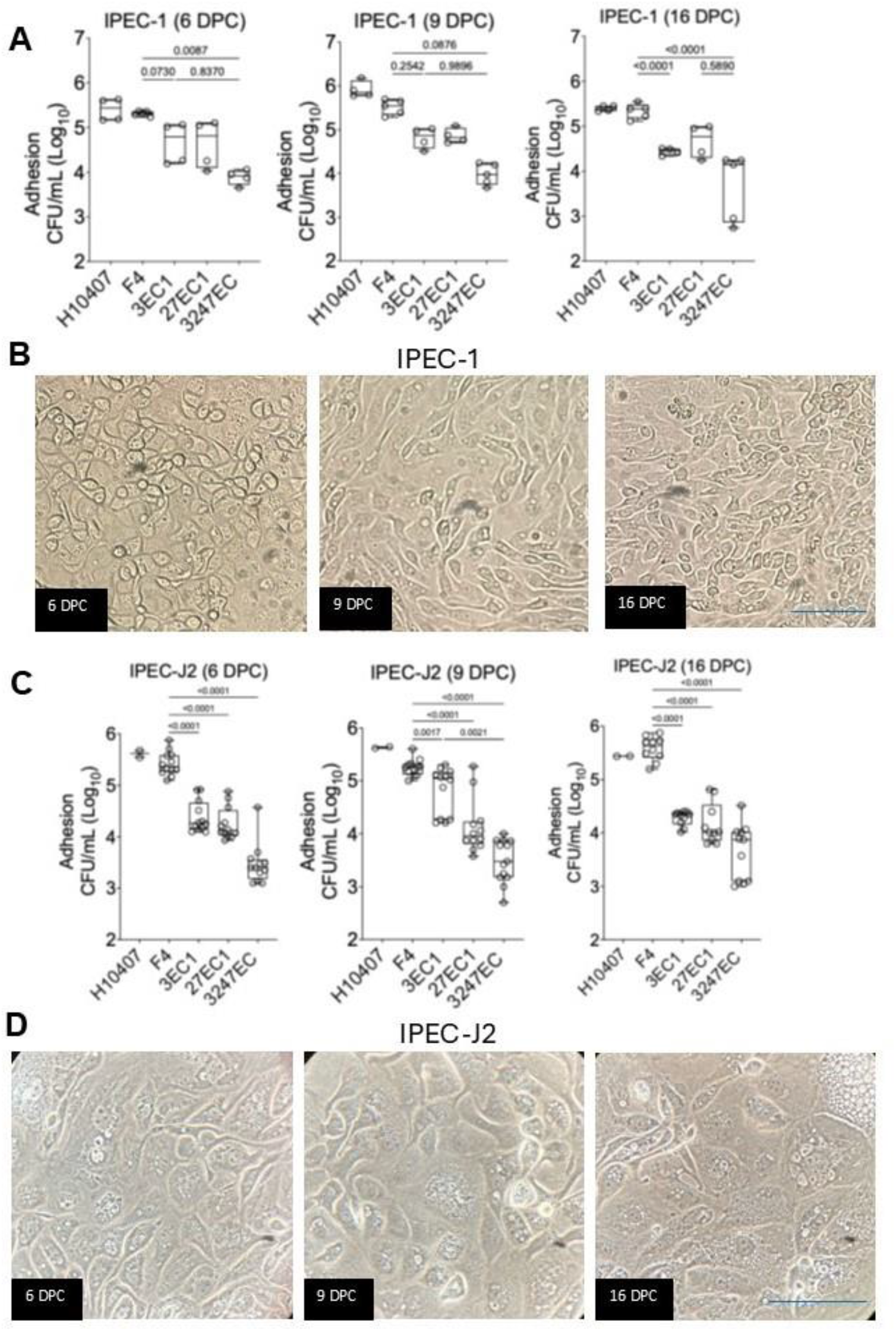
Age-differentiated adhesion comparison of ETEC strains F18 and F4 on porcine intestinal IPEC-1 (A,B) and IPEC-J2 (C,D) cells. **(A,C)** Adhesion characteristics of ETEC F4 and F18 strains to IPEC-1 (**A**) or IPEC-J2 (**C**) cells after 6, 9 and 16 days of post-confluence (DPC), 30 min post-infection at MOI 10. Data represent mean ± SEM. ***, p<0.001; ****, p<0.0001. (**B**,**D**) Photomicrograph of IPEC-1 (**B**) and IPEC-J2 (**D**) after 6,9, and 16 DPC. Scale bar: 50 µm.

Adhesion analysis of four swine ETEC strains revealed distinct maturity-dependent adhesion patterns. In the IPEC-1 model, strain 3EC1 showed a marked increase in adhesion at 9 DPC, reaching levels nearly comparable to those of F4. However, this increase was not observed at 6 or 16 DPC. In contrast, strains 27EC1 and 3247EC demonstrated consistent adhesion capacity across all maturity phases. Strain F4 and H10407 exhibited the highest adhesion level among all tested strains, surpassing the ETEC-F18 strains by an order of magnitude across all maturity phases (**Fig. 2A**). Interestingly, a similar trend of adhesion pattern was observed in the IPEC-J2 model (**Fig. 2B** and **D**).

Since F18 binding to IPEC cells is directly proportional to receptor (FUT1 and FUT2) expression levels (14,15), we analyzed mRNA levels for FUT1 and FUT2 in IPEC cells at 6, 9, and 16 DPC. Compared to IPEC-J2, IPEC-1 showed significantly higher expression of *FUT1* at 6 and 16 DPC and *FUT2* at 16 DPC (**Fig. 3A**). When comparing the two models in parallel, most strains exhibit ∼2-fold stronger adhesion profiles on IPEC-1 than on IPEC-J2, except for 3EC1 at 6 and 9 DPC. Notably, this enhanced adhesion characteristic in IPEC-1 diminished by 16 DPC to a level like that in IPEC-J2. However, at 16 DPC, two of the three F18 strains (3EC1 and 27EC1) show an increasing trend in adhesion capacity (**Fig. 3B**).

**FIG 3.**
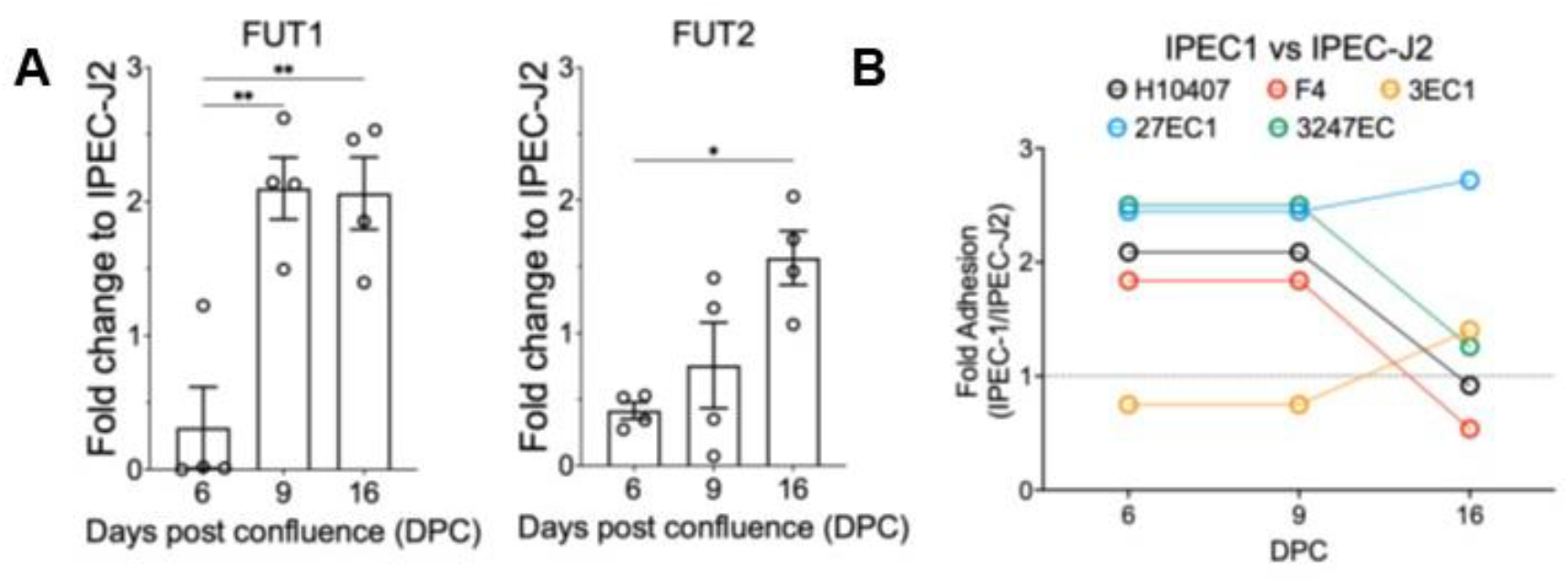
F18-specific host receptor analysis. (**A**) Relative mRNA expression level of FUT1 and FUT2 of IPEC-1 to IPEC-J2 at day 6, 9 and 16 post cell confluence. (**B**) Fold changes of ETEC adhesion on IPEC-1 to IPEC-J2 at day 6, 9 and 16 post cell confluence. Data represent mean ± SEM. *, p<0.05; **, p<0.001.

### Adhesion comparison of pathogenic ETEC strains on porcine intestinal cell lines

Two porcine intestinal epithelium cell lines (IPEC-1 and IPEC-J2) with distinct susceptibility to ETEC fimbriae types were tested for differential ETEC adhesion (21). The IPEC-J2 line is previously documented for its heightened susceptibility to ETEC F4 adhesion, whereas ETEC F18 adheres strongly to IPEC-1 with higher expression of F18 receptors (19). Identical experimental procedures and conditions were applied to both cell lines to ensure compatibility of results. In general, all F18 isolates showed a 1-2 logs reduction in adhesion to both porcine intestinal epithelial cell lines compared to F4 (**Fig. 4A**). Strains 27EC1 and 3247EC adhered markedly less to IPEC-J2 but only slightly reduced to IPEC-1 than the F4 strain. This aligns with the adhesion characteristic of ETEC F18. By contrast, the ETEC F4 and human ETEC H10407 display comparable adhesion to IPEC-1, with H10407 showing slightly higher binding to IPEC-2 (**Fig. 4A**).

**FIG 4.**
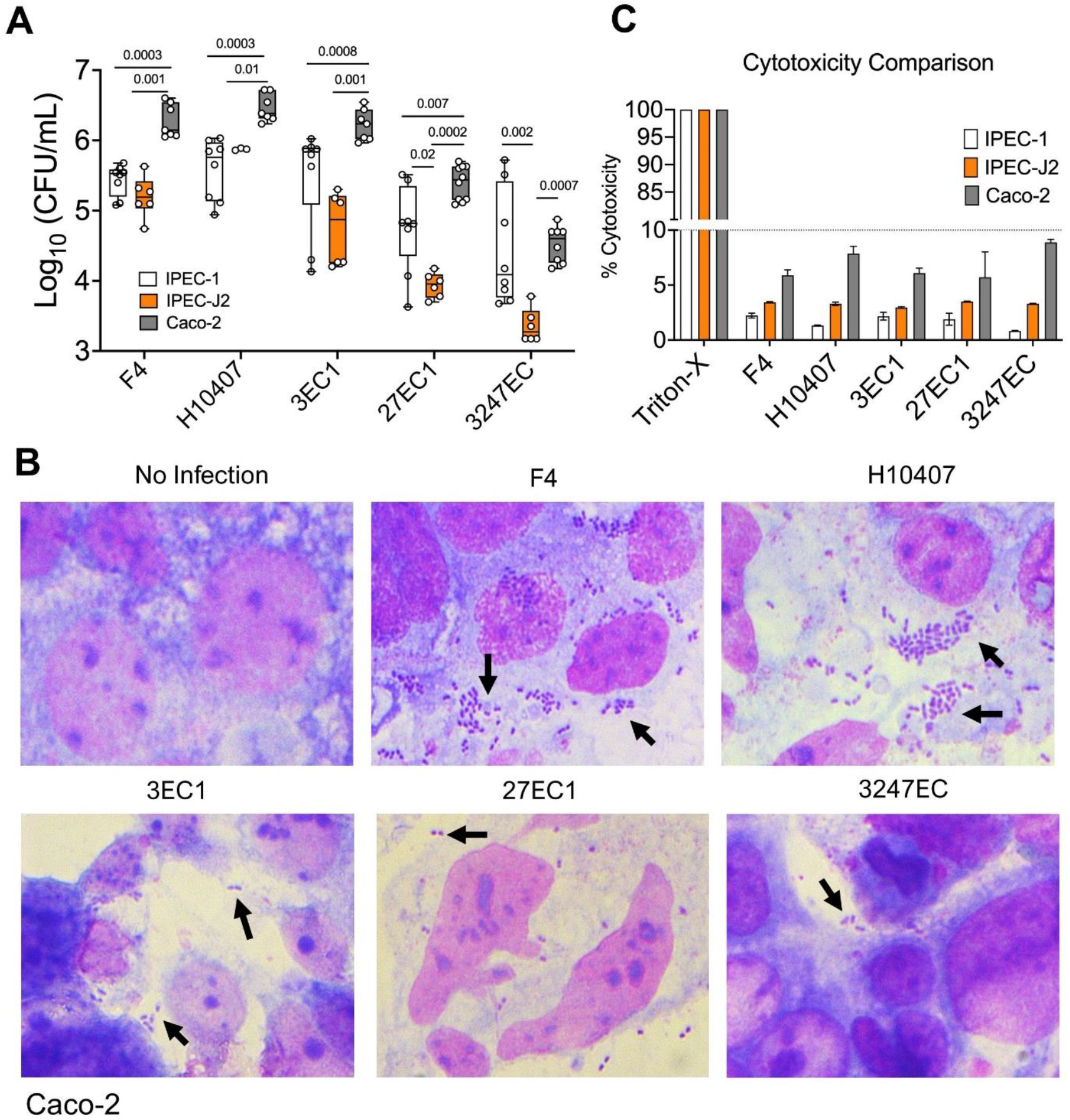
Adhesion comparison of enterotoxigenic *Escherichia coli* (ETEC) strains on porcine intestinal cell lines. (A) Comparative analysis of adhesion of ETEC strains to swine intestinal IPEC-1, IPEC-J2, and Caco-2 cell lines. (**B**) Giemsa staining showing differential adhesion of ETEC strains to Caco-2 cell monolayers. (**C**) Cytotoxicity assay using lactate dehydrogenase (LDH) release from cell monolayers after 30 min of ETEC exposure during the adhesion experiment. Triton-X was used as a positive control. Data are mean ± standard error of the mean (SEM).

These swine-derived ETEC strains were also tested on the human enterocyte-like intestinal Caco-2 cell line as a control. In the Caco-2 cell line, human-origin ETEC H10407 exhibited the highest adhesion as demonstrated by plate counting and Giemsa staining (**Fig. 4A** and **B**). Both 3EC1 and F4 demonstrated relatively high adhesion levels at 5 logs. In contrast, adhesion by 27EC1 and 3247EC was 1-2 logs lower than F4, with 3247EC demonstrating the most significant reduction relative to F4 and H10407.

Post-adhesion lactate dehydrogenase (LDH) levels (% cytotoxicity) in the culture supernatant remained below 10% of the positive control, with no significant alterations observed across all tested groups. These findings indicate that 30 minutes of incubation is sufficient for ETEC to initiate host adhesion but not enough to cause cell damage due to subsequent invasion or toxin release (**Fig. 4C**).

### Genomic analysis of three clinical ETEC F18 isolates

The Whole Genome Sequencing was performed to systematically characterize the virulence gene content of the three clinical F18 isolates with accession numbers: 3EC1 (CP199153), 27EC1 (CP199154), and 3247EC (CP199155) (**Fig. 5, Fig. S2**). The genomes ranged in size from 5 to 5.5 Mbp, with coding sequences (CDS) varying from 5009 to 5701, and an average GC content of 50.7% (**Table 1**). Notably, plasmids were detected only in 3247EC, as summarized in **Table S1**.

**TABLE 1.** General information of *Escherichia coli* strains 3EC1, 27EC1, and 3247EC compared with ETEC F4, H10407, and *E. coli* Nysø.

| Isolate | F4 | H10407 | 3EC1 | 27EC1 | 3247EC | *E. coli* Nysø |
| --- | --- | --- | --- | --- | --- | --- |
| Serogroup | O141:H4 | O78: H11 | O3: H45 | O35: H7 | O119:H23 | O0:H19 |
| MLST | 5786 | 48 | 4214 | 1642 | 224 | 90 |
| FimCHType | 11-560 | 11-398 | 4-31 | 4-31 | 4-0 | 11-41 |
| Genome size (Mbp) | 5.2 | 5.3 | 5.2 | 5.0 | 5.6 | 5.2 |
| GC content (%) | 50.4 | 50.7 | 50.8 | 50.6 | 50.8 | 50.9 |
| Plasmids | IncFIB<br>IncFIC<br>IncFII<br>IncI1-I<br>p0111 | IncFII | ND | ND | Col156<br>IncB/O/K/Z<br>IncFIA<br>IncFIB<br>IncFII(pCo<br>o)<br>IncHI2<br>IncHI2A<br>IncQ1<br>IncY | Col156<br>IncFIB<br>IncFIC<br>IncFII<br>IncI1-I<br>IncX1 |
| Total coding sequence | 5182 | 5322 | 5314 | 5009 | 5701 | 5419 |
| Accession Number | CP002729.1 | FN64941<br>4.1 | This study<br>(CP1991<br>53) | This study<br>(CP1991<br>54) | This study<br>(CP19915<br>5) | DADUQP00000<br>0000 |

**FIG 5.**
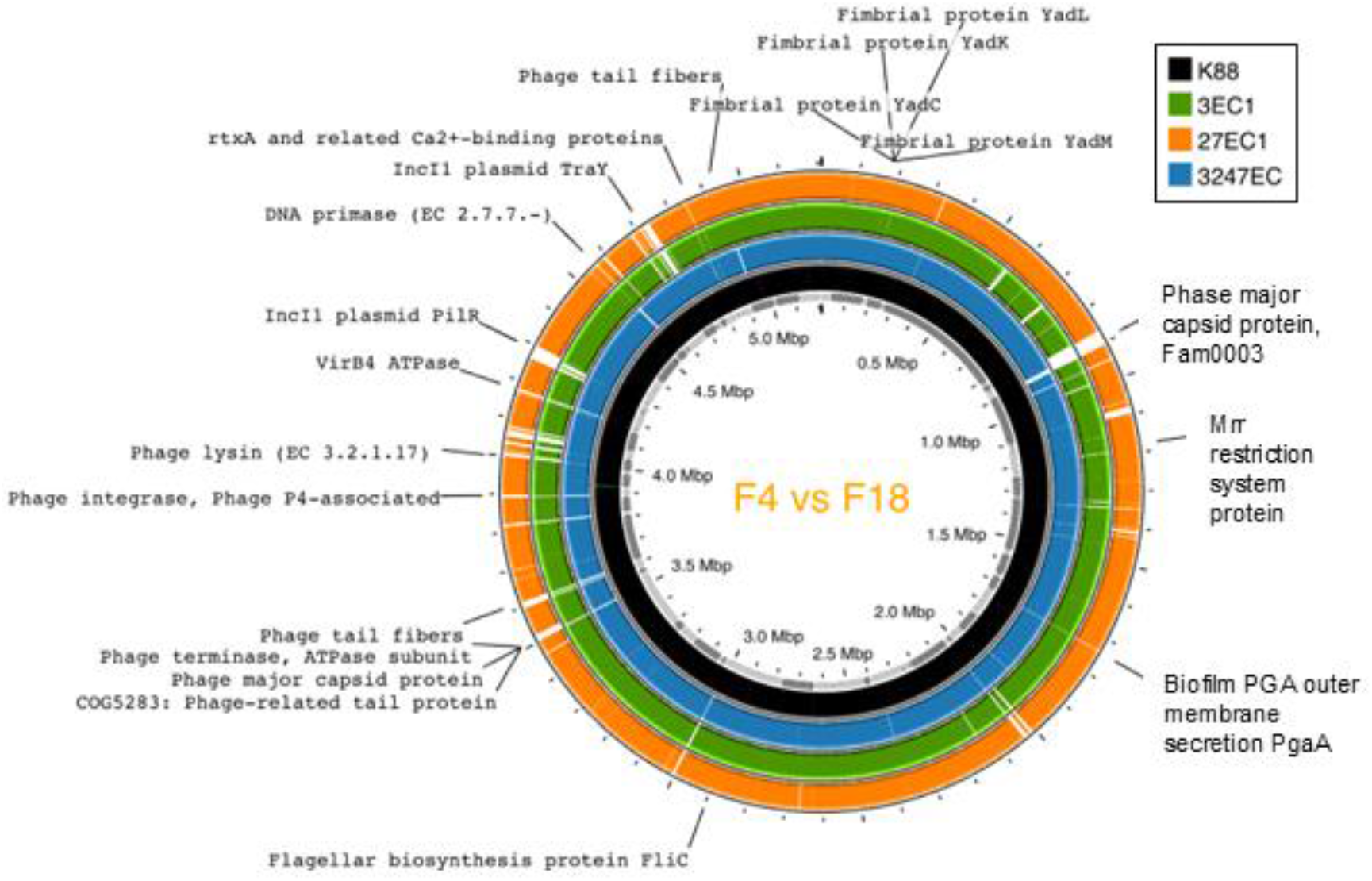
Genome sequence analysis. Genome-wide comparative analysis to examine the genetic differences and similarities between enterotoxigenic *Escherichia coli* (ETEC) F18 isolates and F4 (K88).

Phylogenetic analysis showed that 27EC and 3247EC are more closely related to ETEC F4. At the same time, 3EC1 is genetically distant, clustering closer to the same clade of human ETEC isolate H10407 and ancestral ETEC strain *E. coli* Nysø. For comparison, one historical F18 isolate (Nysø) and one human ETEC (H10407) isolate were included in the analysis (**Fig. 6**).

To assess the virulence potential of these newly isolated F18 strains, ETEC F4 was used as the reference genome for comparative genomics. The analysis identified the top four categories of differential genes as those associated with fimbriae, flagella, LPS synthesis proteins, and phage-associated proteins (**Fig. 5, Fig. S2, Table S2**). The most notable contrasts were observed in virulent genes contributing to pathogen adhesion and toxin synthesis, such as those encoding fimbriae and LPS-associated synthesis genes. Intriguingly, while fimbrial and LPS synthesis genes from 3EC1 showed limited similarity to those from 27EC1 and 3247EC, they closely aligned with F4 and H10407 (**Fig. S2**). For instance, YadU and YadC, key adhesins involved in colonization and biofilm formation (24), exhibited less than 50% homology in 27EC1 and 3247EC compared to 3EC1 and F4. In contrast, flagellar-associated genes were found to be relatively conserved, showing over 80% amino acid similarity among all swine- and human-derived ETEC strains (**Fig. 6A, Fig. S2**).

**FIG 6.**
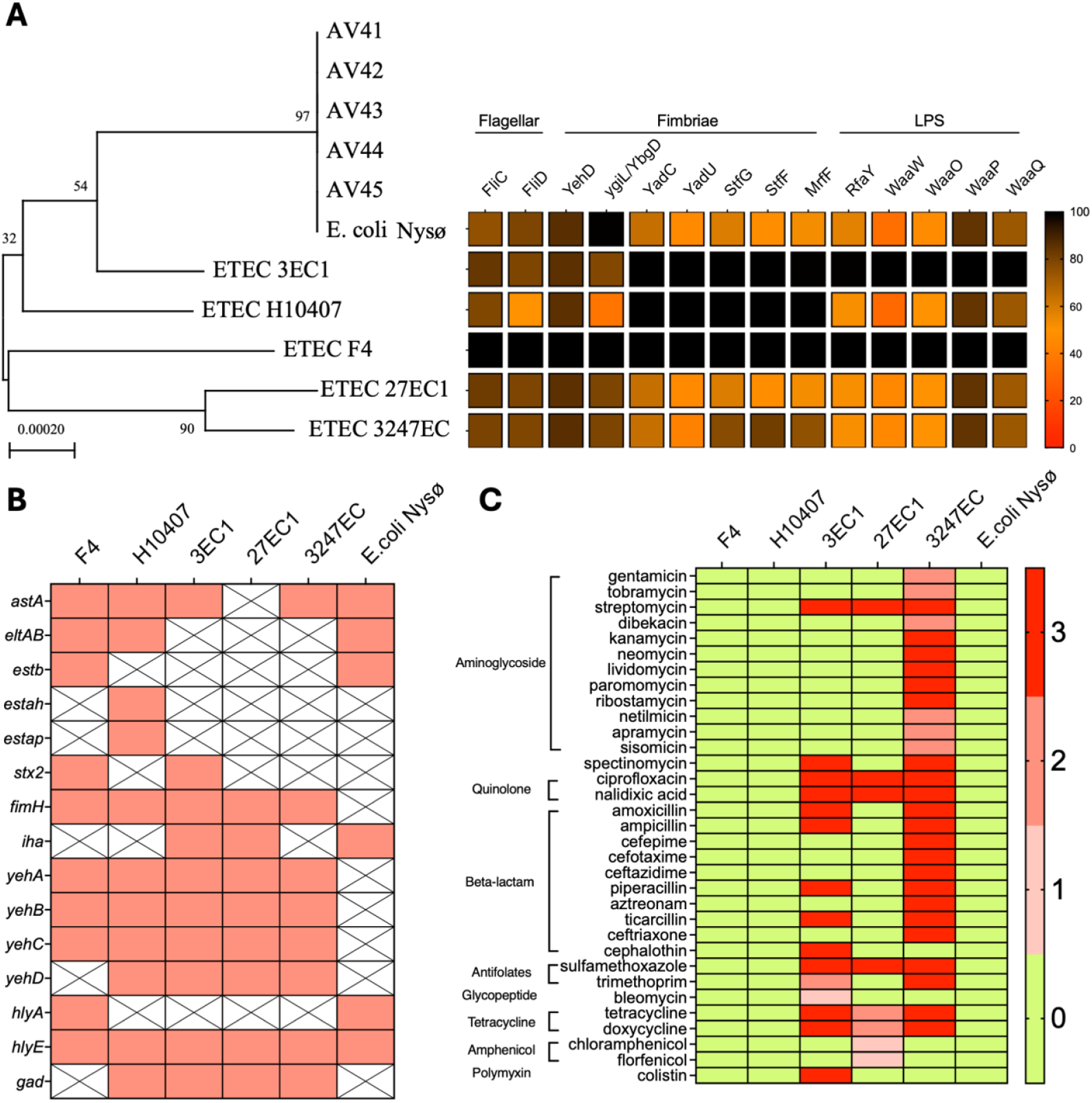
Comparative genetics of human and swine pathogenic *Escherichia coli* strains. (**A**) Phylogenetic tree and similarity of core virulent genes of enterotoxigenic *Escherichia coli* (ETEC) clinical isolates from swine and human. The phylogenetic tree was constructed based on core conserved genes, root on midpoint. Bootstrap value is 1000. For comparative virulent gene analysis, a scale from 0 to 100 reflects the similarity of each gene using ETEC F4 as a reference. (**B**) Comparative genomics showing the presence and absence of virulent genes. (**C**) Prediction of antibiotic resistance phenotype. Value code: 0, No match found; 1: Match < 100% ID and match length < ref length; 2: Match = 100% ID and match length < ref length; 3: Match = 100% ID and match length = ref length.

Classic ETEC toxin genes encoding heat-labile (*eltAB*) or heat-stable (*esta, estb*) toxins were absent in all three ETEC F18 strains. However, non-classic ETEC toxins, such as EAST-1, were identified with minimal variation between 3EC1 and 3247EC. Interestingly, *stx2e*, a Shiga toxin gene, was identified in 3EC1, but not in 27EC1 and 3247EC, indicating potential hybrid virulent traits. The avian hemolysin (*hlyE*) (25) was presented in all ETEC F18 and H10407, which aligned with hemolytic activity (**Fig. 1**). While both hemolysin A and avian hemolysin were detected in F4 and *E. coli* Nysø. Outer membrane fimbrial cluster Yeh family proteins (YehA-D) were identified in both human and porcine ETECs, except *E. coli* Nysø. Additionally, glutamate decarboxylase (GAD), an enzyme that aids in acid resistance (e.g., stomach fluid), was detected in H10407, 3EC1, 27EC, and 3247EC. All ETEC strains harbored the *fimH* fimbrial genes, except *E. coli* Nysø. The adhesin *iha*, prevalent in pathogenic *E. coli* (26), was identified in 3EC1, 27EC1, and *E. coli* Nysø (**Fig. 6B**). A comprehensive list of virulence factors is summarized in **Table S2**.

### Antibiotic resistance phenotype

Genomic analysis revealed that all three clinical ETEC F18 isolates carry several antibiotic resistance-associated genes (**Fig. 6C, Table S3**), which are absent in the ETEC F4 and H10407. Interestingly, the isolates harbor genes conferring resistance to aminoglycosides, aminocyclitols, quinolones, beta-lactams, folate pathway antagonists, and tetracyclines. Antibiotic genes conferring resistance to streptomycin, ciprofloxacin, nalidixic acid, and sulfamethoxazole are identified in all three ETEC isolates.

We phenotypically verified the antibiotic resistance genotype using microdilution (**Table 2**) and disc diffusion (**Table 3**) assays against a panel of antibiotics. Phenotypic results confirmed the resistance in the three clinical F18 isolates. Genomic prediction of strains 3EC1 and 27EC1 were resistant to penicillin (amoxicillin, ampicillin, ticarcillin, and piperacillin), aminoglycosides (gentamicin, kanamycin, neomycin, netilmicin, paromomycin, apramycin, sisomicin, streptomycin, tobramycin), aminocyclitol (closely related to aminoglycosides, spectinomycin), cephalothin, sulfamethoxazole, and florfenicol. However, 3247EC was found to be sensitive to β-lactam antibiotics, including penicillin derivatives (amoxicillin, ampicillin, ticarcillin, and piperacillin), as well as aminoglycosides (gentamicin, kanamycin, neomycin, netilmicin, paromomycin, apramycin, sisomicin, streptomycin, and tobramycin), which contrasts with the antimicrobial resistance (AMR) prediction. Interestingly, we observed a morphological change in 3247EC upon ampicillin treatment, characterized by bacterial elongation. This atypical morphology reverted to the typical rod shape upon removal of ampicillin. In contrast, no such morphological changes were observed in 27EC or 3EC1 under the same treatment conditions (**Fig. S3**).

**TABLE 2.** Minimum inhibitory concentrations (MIC, µg/mL) of select antibiotics against *Escherichia coli* strains 3EC1, 27EC1, 3247EC, H10407, and F4(K88).

|  | *Escherichia coli* strain |  |  |  |  |
| --- | --- | --- | --- | --- | --- |
|  | 3EC1 | 3247EC | 27EC1 | H10407 | F4(K88) |
| **Amoxicillin** | >128 | 4 | >128 | 2 | 8 |
| **Ampicillin** | >128 | 4 | >128 | 2 | 16 |
| **Apramycin** | >128 | 64 | >128 | 32 | 16 |
| **Aztreonam** | 0.25 | 0.12 | 0.12 | 0.03 | 0.12 |
| **Bleomycin** | 2 | 1 | 1 | 2 | 1 |
| **Cefepime** | 0.12 | 0.06 | 0.06 | 0.03 | 0.03 |
| **Cefotaxime** | 0.06 | 0.06 | 0.06 | 0.03 | 0.06 |
| **Ceftazidime** | 0.25 | 0.25 | 0.5 | 0.12 | 0.25 |
| **Ceftriaxone** | 0.12 | 0.12 | 0.12 | 0.06 | 0.12 |
| **Cephalothin** | 32 | 32 | 32 | 8 | 32 |
| **Chloramphenicol** | 2 | 2 | 2 | 0.5 | 2 |
| **Colistin** | 0.25 | 0.25 | 0.25 | 0.25 | 0.25 |
| **Doxycycline** | 16 | 8 | 32 | 0.5 | 2 |
| **Florfenicol** | 8 | 8 | 8 | 2 | 4 |
| **Gentamicin** | >128 | 1 | 32 | 1 | 0.5 |
| **Kanamycin** | >128 | 4 | >128 | 8 | 2 |
| **Neomycin** | >128 | 2 | 128 | 2 | 2 |
| **Netilmicin** | >128 | 2 | 32 | 1 | 0.5 |
| **Paromomycin** | >128 | 4 | >128 | 4 | 2 |
| **Piperacillin** | >128 | 2 | >128 | 1 | 2 |
| **Sisomicin** | >128 | 1 | 16 | 2 | 0.5 |
| **Spectinomycin** | >128 | 32 | >128 | 32 | 16 |
| **Streptomycin** | >128 | >128 | >128 | 8 | 2 |
| **Sulfamethoxazole** | >512 | >512 | >512 | >512 | 16 |
| **Tetracycline** | 32 | 128 | >128 | 2 | 2 |
| **Ticarcillin** | >128 | 16 | >128 | 8 | 16 |
| **Tobramycin** | >128 | 1 | 32 | 1 | 0.5 |
| **Trimethoprim** | >128 | >128 | 0.12 | 0.12 | 0.25 |

**TABLE 3:** Inhibition zones (mm) of the antibiotics against *Escherichia coli* strains after agar disc diffusion assay.

| **Antibiotics** | ***Escherichia coli* strain** |  |  |  |  |  |
| --- | --- | --- | --- | --- | --- | --- |
|  | 3EC1 | 3247EC | 27EC1 | H10407 | F4(K88) | 3247EC+Amp |
| Ampicillin | 0 | 21 | 0 | 23 | 22 | 25 |
| Sulfamethoxazole | 0 | 0 | 28 | 24 | 27 | NT |
| Clindamycin | 0 | 0 | 8 | 0 | 0 | NT |
| Penicillin G | 0 | 0 | 0 | 11 | 9 | NT |
| Gentamicin | 0 | 19 | 10 | 18.5 | 18 | 18.2 |
| Tetracycline | 23 | 8 | 0 | 30 | 25 | 7 |
| Vancomycin | 0 | 9 | 0 | 9 | 10 | 8 |
| Cephalothin | 17 | 17 | 17 | 24 | 18 | 22 |
| Ciprofloxacin | 0 | 25 | 27 | 35 | 33 | 21.6 |
| Chloramphenicol | 24 | 23 | 29 | 33 | 25 | 24 |

NT= Not tested

## Discussion

The swine industry suffers from significant economic losses due to piglet mortality from ETEC strains (8,27). Yet molecular information, antibiotic resistance, and virulence phenotype data are limited (28,29). In this study, we conducted a comprehensive analysis of three clinical *E. coli* isolates from U.S. pig farms, elucidating their respective pathobiology through molecular & cellular assays and whole-genome sequencing. Notably, we identified the *E. coli* isolates adhesion efficiency to swine intestinal cells peaks at 9 days post-confluence, coinciding with the progressive upregulation of the F18 receptor (F18R/FUT1) in the neonatal porcine intestine during the suckling phase (30). This temporal correlation underscores the feature of ETEC F18 infection. The lack of significant differences in F18 ETEC adhesion between the IPEC-1 and IPEC-J2 cell lines may be a threshold effect in gene expression. The observed differences in FUT1/2 expression levels are too subtle to manifest as a functionally distinct adhesion profile. Specifically, ETEC F4 still outcompetes F18 strains in the IPEC-1 model, suggesting that the cell lines express a substantial level of F4-specific receptors as well. Ultimately, these results underscore that ETEC adhesion is a complex multifactorial process determined by factors beyond FUT1/2 expression levels alone.

Among the three F18 isolates, 3EC1 exhibited the most pronounced virulent phenotype. Phylogenetic analysis shows a closer evolutionary trajectory between porcine ETEC 3EC1 and human ETEC H10407, suggesting potential cross-species transmission or shared evolutionary pathways. Comparative genomic and phylogenetic analysis revealed that 3EC1 harbors genetic features resembling both porcine F4-like and human ETEC pathovars, particularly in fimbrial biosynthesis and LPS synthesis pathways. These traits were functionally validated across porcine and human intestinal cell line models. Furthermore, WGS also detected *stx2e*, a defining toxin of Shiga toxin-producing *E. coli* (STEC), in F4 and 3EC1, consistent with the prevalence of swine ETEC isolates in the U.S. (31) and China (32). This finding suggests the emergence of novel *E. coli* hybrid pathovars with combined virulence mechanisms, potentially complicating clinical management and zoonotic risk.

Another critical concern arising from this study is the pervasive antibiotic resistance observed in these isolates, which contrasts with the resistance profiles of classical ETEC F4 and F18 serovars. Both 3EC1 and 3247EC exhibit extended-spectrum β-lactamase (ESBL) phenotypes by carrying bla_CTX-M-15_ and bla_TEM_, respectively. Resistance to ampicillin is the most commonly observed trait in *E. coli* isolates from animal farms in Europe and America in recent years (33). A clinical survey of *E. coli* isolates from Bangladesh reported bla_CTX-M-15_ as the most prevalent β-lactamase antibiotic-resistant gene, detected in 52% of cases, while bla_TEM_ was identified in 20% of cases (34). Surprisingly, no antibiotic resistance phenotypes such as ampicillin and gentamicin were observed during phenotypic testing, despite the presence of resistance genes. Notably, 3247EC is the only isolate found to carry plasmids. The loss of resistance in 3247EC may be due to plasmid loss during sub-culturing (**Table 4**), which may be associated with changes in morphology (**Fig. S3**). The dissemination of ESBL is driven mainly via horizontal gene transfer (HGT) from plasmid-borne elements (29).

**TABLE 4.**
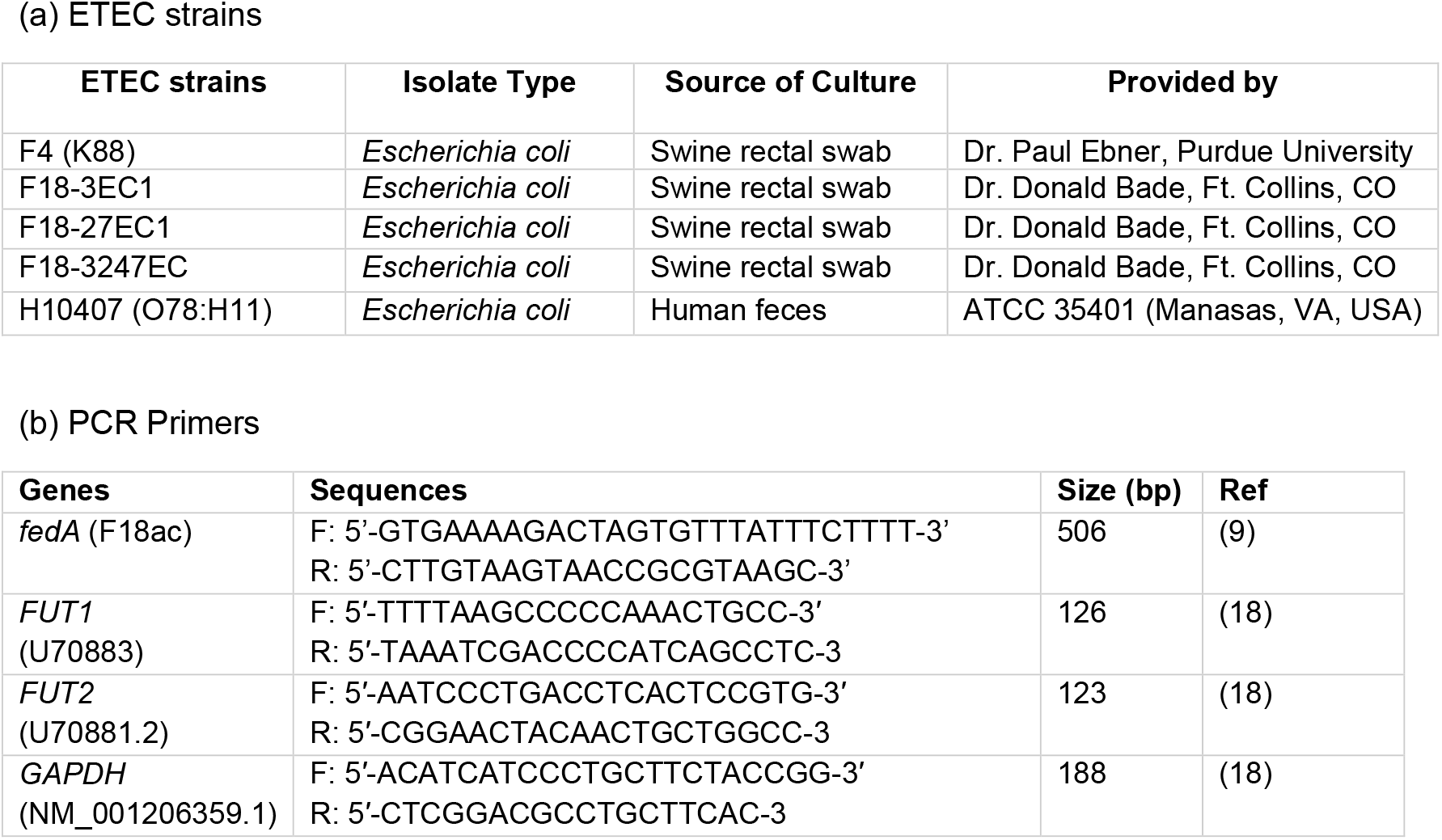
Enterotoxigenic *Escherichia coli* (ETEC) strains (a) and primers (b) used in the study.

Additionally, 3EC1 and 27EC1 exhibited elevated resistance to aminoglycosides, consistent with previously reported patterns in ETEC F18 (29). This may be attributed to the widespread use of gentamicin in the treatment of neonatal colibacillosis in pig farms (35). The ETEC F18 isolates from animal farms have shown significantly increased resistance to gentamicin and kanamycin; a trend is not observed in ETEC F4 (33). The increasing prevalence of multidrug-resistant ETEC isolates poses a significant public health threat, as HGT facilitates the rapid spread of resistance determinants across microbial communities. Given the diversity and adaptability of these communities, continuous genetic monitoring and surveillance are essential to track emerging resistance patterns and inform mitigation strategies. Collectively, these findings emphasize the dynamic evolution of ETEC pathovars, driven by both virulence gene diversification and antibiotic selection pressures. To mitigate the risks of emergent hybrid strains and multidrug resistance, sustained genomic surveillance and stringent antibiotic stewardship in livestock systems are imperative. Such measures will safeguard animal and human health against these adaptable pathogens.

## MATERIALS AND METHODS

### Bacterial strains and growth conditions

Porcine ETEC clinical strains (F4, 3EC1, 27EC1, 3247EC) were initially isolated from swine rectal swabs, while the human reference strain H10407 (ATCC35401), was isolated from human feces (**Table 4**). Bacterial cultures were inoculated from single colonies and grown in tryptic soy broth (30 g/L) supplemented with yeast extract (6 g/L) (TSBYE) (BD BBL, Franklin Lakes, NJ, USA) in an orbital shaker at 150 rpm, 37°C for 12–16 h. Freshly prepared stationary-phase cultures were used in all experiments, with an OD_600_ of approximately 1.5. *L. monocytogenes* F4244 was grown in TSBYE at 37°C for 12–16 h.

### DNA isolation and PCR determination of virulence factor in ETEC strains

DNA was extracted using the Zymo Research Quick-DNA Fungal/Bacterial Kits according to the manufacturer’s instructions. The quality (A260/280 ∼1.8) and purity (A260/230 > 2.0) of the extracted DNA were assessed using a NanoDrop spectrophotometer (Thermo Fisher Scientific, NJ, USA) and agarose gel electrophoresis. For PCR analysis (9), approx. 10 ng of genomic DNA (gDNA) served as the template for amplification of the *fedA* gene (**Table 4b**).

Thermocycling conditions include initial denaturation at 95°C (30s); 30 cycles of 95°C (15s), 50°C (15s), 68°C (30s) with a final extension at 68°C (120s). The amplified products were analyzed by agarose gel electrophoresis and visualized under UV light at 320 nm.

### Reverse-transcriptase quantitative PCR for the expression F18 receptors

Swine intestinal epithelial cell lines (IPEC-1 and IPEC-J2) were maintained at 37°C under 5% CO_2_ in RPMI-1640 medium supplemented with 10% FBS. Upon reaching confluence, cells were cultured for an additional 6, 9 and 16 days prior to total RNA extraction. RNA quality is ensured (A280/260 > 2.0) and normalized to 2000 ng before reverse transcription (NEB). Transcribed cDNA (NEB) was 10-fold diluted (the total reverse transcription mixture was 20 µL, diluted to 200 µL by adding 180 µL DEPC water, and 3 µL of cDNA from each sample was used for PCR) in ultra-pure water (ThermoFisher). The diluted cDNA was used as a template for qPCR amplification for *FUT1 and FUT2*, as before (18). *GAPDH* was used as an internal reference gene. The RT-qPCR primers used are listed in **Table 4b**. 2^-ΔΔCT^ method was used to calculate the relative expression of the *FUT1/2* in IPEC-1 and IPEC-J2 cell lines.

### Whole genome sequence analysis and genotyping

Bacterial genome sequencing was outsourced to GenScript (Piscataway, NJ, USA) for library preparation and sequencing on the Illumina NextSeq platform using 150 bp paired end reads. Raw reads were assembled using Shovill (Galaxy v1.0.4), and the assembly quality was evaluated using QUAST (36). The assembled genomes of ETEC isolates were annotated and comparatively analyzed using the RAST server (37). Plasmids, virulence factors, antibiotic resistance genes, fimbrial type, serotype (O and H serogroups), and multilocus sequence types (MLST) are identified by phenotyping services from the Center for Genomic Epidemiology (https://www.genomicepidemiology.org/services/).

### Genome annotation, phylogenetic analysis, and comparative genetics

In this study, 11 bacterial genomes were annotated using GTDB-Tk (38). A core conserved gene approach was utilized to identify and extract the gene sequences of 120 conserved proteins from the genome data. The resulting protein sequences were aligned through multiple sequence alignments. Phylogenetic analysis was subsequently conducted on the aligned sequences of 120 core genes using MEGA 11 software (39), employing the Neighbor-Joining (NJ) method with 1000 bootstrap replicates.

### Bacterial adhesion to porcine and human intestinal cell lines

To further validate the differential ETEC adhesion, two porcine intestinal epithelium cell lines (IPEC-1 and IPEC-J2) with distinct susceptibility to ETEC fimbriae types were used. The IPEC-J2 cell line is a morphologically and metabolically more active cell model with higher expression of microvilli than the IPEC1 cell line (21). IPEC-1 and IPEC-J2 purchased from Leibniz Institute DSMZ (Braunschweig, Germany) were cultured in complete RPMI-1640 medium (Thermo Fisher Scientific) supplemented with 4 mM L-glutamine, 1 mM sodium pyruvate, 10% fetal bovine serum (FBS, Atlanta Biologicals), and 0.0001% epidermal growth factor (EGF, Corning Life Sciences). The human colon-derived Caco-2 cells were cultured in complete Dulbecco’s Modified Eagle’s Medium (DMEM) containing 10% FBS. To investigate the adhesion capacity of ETEC, IPEC-1, IPEC-J2 cells at three maturation stages (6, 9, and 16 days post-confluence), designated as “Early,” “Mid,” and “Late,” were used as a model to mimic the natural physiology of the intestine and assess corresponding adhesion dynamics (**Fig. S2A, C**).

For adhesion assays, stationary-phase bacterial cultures were diluted to a multiplicity of infection (MOI) of 10 (5×10^6^ to 1×10^7^ CFU/well according to cell density) in serum-free RPMI-1640, unless otherwise stated. Bacterial cultures were centrifuged at 6000× g for 5 min, washed with sterile phosphate-buffered saline (PBS), and resuspended in 500 µL of serum-free RPMI-1640. To facilitate bacterial contact with the cell layers, 0.5 mL of the bacterial suspension was added to each well of the 12-well plates, ensuring minimal liquid volume. The cells were incubated at 37°C for 30 min to allow bacterial adherence. After incubation, non-adherent bacteria were removed by washing the cells with serum-free RPMI-1640. To quantify adhered bacteria, the cells were lysed with 0.1% Triton X-100 in serum-free RPMI-1640. This concentration effectively lysed mammalian cells without affecting bacterial viability, releasing any intracellular bacteria. The number of adherent bacteria was determined by serial dilution and plating on MacConkey agar (BD BBL, Franklin Lakes, NJ, USA). Each experiment was conducted in duplicate.

### ETEC-induced cytotoxicity

To evaluate the cytotoxic effects of ETEC, if any, on three different cell models during the adhesion assay, the supernatants were collected after bacterial infection and analyzed for lactate dehydrogenase (LDH) activity using the LDH Cytotoxicity Assay Kit (Cayman Chemical Company, Ann Arbor, MI). The LDH assay, an indicator of cell damage, quantifies the release of LDH from compromised cells into the culture media, providing a measure of the percentage of cytotoxicity (40). Supernatants from untreated cells served as a negative control, while a solution of 1% Triton X-100 in RPMI-1640, which fully lyses cells to release maximal LDH, was used as a positive control. Each experimental condition was performed in triplicate. Absorbance was measured spectrophotometrically at 490 nm using a microplate reader. The percentage of cytotoxicity was then calculated (40)

### Wright-Giemsa staining

Wright-Giemsa staining was employed to visualize infected cellular structures and the binding of bacterial pathogens to cells. Cell lines were grown in chambered Lab-Tek™ II slide flaskets (Thermo Scientific) to about 80% confluence. Following bacterial infection (MOI 10) for 1 h at 37ºC, the cells were rinsed twice with PBS and fixed with methanol for 5 min. The cells were then flooded with Giemsa stain solution (10% Giemsa stain, 10% methanol, and 80% PBS) for 45 min, rinsed with PBS for 1 min, and then rinsed with deionized (DI) water. The slides were air-dried and examined under a Leica light microscope (Wetzlar, Germany).

### Microdilution assay to test antibiotic resistance

Antibiotics were dissolved in water or dimethyl sulfoxide (DMSO) at a concentration of 10 - 40 mg/mL and were visually checked to ensure complete dissolution. *E. coli* strains were streaked on tryptic soy agar (TSA) plates and incubated aerobically at 37ºC overnight. As recommended by the Clinical and Laboratory Standards Institute (41), colonies were suspended in PBS and diluted in Mueller-Hinton broth to achieve a concentration of 5 ×10^5^ CFU/mL. The bacterial suspensions were then mixed with 2-fold serial dilutions of the antibiotics in 96-well plates (in duplicates). Colistin was used as a positive control, while DMSO was used as a negative control. Plates were incubated aerobically at 37ºC for 16 h before inspected for bacterial growth. The minimum inhibitory concentration (MIC) of each antibiotic was the lowest concentration at which the wells were clear, indicating no bacterial growth.

### Agar disk diffusion assay

Bacterial isolates were grown overnight on TSA plates at 37ºC. A bacterial suspension of 1 x 10^8^ CFU/mL was prepared in PBS and was spread evenly on the surface of Mueller-Hinton agar plates. Antimicrobial susceptibility discs (Oxoid, Lenexa, KS) were aseptically placed on the surface of each inoculated plate and incubated at 37ºC for 16 h. The diameter of the inhibition zone around each disc was measured three times, and the average diameter was reported (**Table 3**). The test was repeated for strain 3247EC on an ampicillin-containing plate (100 µg/mL) to evaluate the effect of including ampicillin on the bacterial resistance pattern.

### Statistical analysis

Experimental data were analyzed using GraphPad Prism 9 (La Jolla, CA). Statistical comparisons between treatments were conducted using one-way or two-way analysis of variance (ANOVA) followed by Tukey’s multiple-comparison test for more than two treatments or Student’s *t*-test for comparisons involving only two treatments. Unless otherwise specified, data from all experiments are presented as the mean ± standard error of the mean (SEM) from two independent experiments.

## ACKNOWLEDGMENTS

This research was partly supported by United Animal Health, Inc. (Sheridan, IN, USA) and the USDA National Institute of Food and Agriculture (Hatch accession no. 1016249). Any opinions, findings, conclusions, or recommendations expressed in this publication are those of the author(s) and do not necessarily reflect the view of the U.S. Department of Agriculture. We thank Dr. Donald Bade and Dr. Paul Ebner for their generosity in providing the ETEC strains.

## DATA AVAILABILITY

Genome sequence information has been deposited at NCBI (PRJNA1311150), and all other data are presented in the manuscript.

## SUPPLEMENTAL MATERIALS

**Supplemental Figures S1 to S3 and supplemental Tables S1 to S3**

**Figure S1. (A)** Growth of F18 clinical isolates on MacConkey agar plates.

**Figure S2.** Genome sequence of enterotoxigenic *E. coli* (ETEC) F18 strains. Genomic map of ETEC (**A**) 3EC1, (**B**) 27EC1, and (**C**) 3247EC displaying the differential presence of genes encoding fimbriae or fimbriae-like proteins (box). Panel D shows a comparative analysis of major virulence factors among ETEC F18 and ETEC F4 strains. Differences in genes encoding flagella, fimbrae, and LPS were seen among F18 strains, while no difference in toxin genes were observed.

**Figure S3.** Morphological characteristics of enterotoxigenic *Escherichia coli* (ETEC) cells after growth in ampicillin-containing or ampicillin-deficient tryptic soy broth (TSB) or Mueller-Hinton agar plates. Magnification 1000x.

**Table S1.** List of plasmids identified in *Escherichia coli* strain 3247EC via PlasmidFinder.

**Table S2.** List of virulent genes identified in *Escherichia coli* strains 3EC1, 27EC1, and 3247EC via VirulenceFinder.

**Table S3.** List of acquired antimicrobial resistance genes in *Escherichia coli* strains 3EC1, 27EC1, and 3247EC.

